# Unveiling cryptic diversity among Müllerian co-mimics: insights from the Western Palaearctic *Syntomis* moths (Lepidoptera: Erebidae)

**DOI:** 10.1101/794602

**Authors:** Andrea Chiocchio, Paola Arduino, Rossella Cianchi, Daniele Canestrelli, Alberto Zilli

## Abstract

Accurate species delimitation is of primary importance in biodiversity assessments and in reconstructing patterns and processes in the diversification of life. However, the discovery of cryptic species in virtually all taxonomic groups unveiled major gaps in our knowledge of biodiversity. Mimicry complexes are good candidates to source for cryptic species. Indeed, members of mimicry complexes undergo selective pressures on their habitus, which results in strong resemblance between both distantly and closely related species. In this study, we used a multi-locus genetic approach to investigate the presence of cryptic diversity within a group of mimetic day-flying moths whose systematics has long been controversial, the Euro-Anatolian *Syntomis*. Results showed incongruence between species boundaries and currently accepted taxonomy of the group. Both mitochondrial and nuclear markers indicate presence of four, well-distinct genetic lineages. The genetic distance and time of divergence between the Balkan and Italian populations of *S. marjana* are the same as those found between *S. phegea* and *S. ragazzii*, the last two being well-distinct, broadly sympatrically occurring species. The divergence between the two lineages of *S. marjana* dates back to the Early Pleistocene, which coincided with substantial changes in climatic conditions and vegetation cover in Southern Europe that have likely induced geographic and ecological vicariance. Our results show that the species richness of mimicry complexes inhabiting temperate regions might still be severely underestimated. *Syntomis* populations up to now designated as S. *marjana albionica*, *S. m. quercii* and *S. marjana kruegeri* s. str. are here considered to represent a separate species from nominate *marjana* and are distinguished as *Syntomis quercii* Verity, 1914, **bona sp.**, **stat. nov**.

## Introduction

Accurate species delimitation is of primary importance for understanding tempo and modes of species diversification (Wiens, 2007). Historically, new species have mainly been recognized using morphological trait variation (Wiens & Servedio, 2000). However, species can evolve even in absence of conspicuous morphological trait divergence, as shown by the discovery of the so-called cryptic species (Bickford *et al.*, 2007). These had originally been spotted on the basis of subtle ecological or behavioural features (Mayr, 1963), though the use of genetic markers to circumscribe biological species has nowadays boosted their discovery (Ayala & Powell, 1972; Knowlton, 1993; Beheregaray & Caccone, 2007; Carstens *et al.*, 2013). The recognition of cryptic species virtually across all taxonomic groups (Pfenninger & Schwenk, 2007), included many well-studied ones (Roca *et al.*, 2001; Fennessy *et al.*, 2016), unveiled major gaps in our knowledge of biological diversity (Beheregaray & Caccone, 2007; Struck *et al.*, 2018). These gaps in turn constrain our understanding of the mechanisms involved in biological diversification and in the establishment of interactions at the community level, and limit the deployment of measures in many areas of biodiversity management, from conservation biology to pest or disease control (Beheregaray & Caccone, 2007; Balint *et al.*, 2011; Robuchon *et al.*, 2019).

Although cryptic species are widespread throughout the tree of life, they are over-reported in some groups, such as freshwater fishes, deep-sea clams, polychaetes, frogs, mites, parasitic insects and nematodes (reviewed in Perez-Ponce & Poulin, 2016). Cryptic species seem to be commoner in animals using non-visual mating signals (e.g. many frogs) and/or under selection promoting morphological stasis or convergent evolution (e.g. parasites) (Bickford *et al.*, 2007; Struck *et al.*, 2018). They are also common in cases of recent and/or rapid speciation, when short divergence time did not allow the accumulation of detectable phenotypic differences (Reidenbach *et al.*, 2012; Gustafsson *et al.*, 2014). However, numerous taxa characterized by life history traits likely encouraging cryptic diversification have been poorly investigated, and many of them are still analysed with a traditional morphological approach (Bickford *et al.*, 2007; Struck *et al.*, 2018). As a consequence, a clear picture of the distribution of cryptic species across the tree of life is still lacking.

Mimicry complexes are good candidate sources of cryptic species. In particular, cryptic species are expected among groups of closely related species involved a same Müllerian mimicry ring (Ruxtoun *et al.*, 2004), or of look-alikes in masquerade rings (cf. Boppré *et al.*, 2017), as phenotypic divergence after speciation is constrained by stabilizing selection on the shared warning signals (but see Lawrence *et al.*, 2019). Interestingly, it was shown that look-alikeness among co-mimics can also be achieved secondarily via introgressive hybridization and incorporation of pattern-determining genes from related species (Giraldo *et al.*, 2008; Pardo-Diaz *et al.*, 2012; Jiggins, 2017). The imbalance between species divergence and phenotypic divergence is a prerequisite for the evolution of cryptic species (Struck *et al.*, 2018). Thus, species involved in mimicry complexes provide a good testing ground to study the contrasting effects of divergence and stasis in the evolution of cryptic species. However, the systematics of these species often results still inaccurate or partial, especially if exclusively based on external morphology. As a matter of fact, the presence of cryptic species within mimicry complexes has been reported almost exclusively for some well-studied tropical taxa, which provided excellent insights on how mimicry can affect patterns of diversification and speciation (Pfennig, 2012, and references therein; Jiggins, 2017). Yet mimicry complexes are not limited to the tropical regions.

In this study, we investigated the presence of cryptic species within a group of mimetic moths occurring in temperate regions whose systematics has long been controversial, the Euro-Anatolian *Syntomis*. These are distasteful, day-flying moths showing aposematic colorations which belong to a Müllerian mimicry complex including also several unrelated co-mimics, e.g. yellow morph of *Zygaena ephialtes*, *Z. transalpina* (Zygaenidae) and possibly *Callimorpha dominula* (Erebidae) (Bullini *et al.*, 1969; Sbordoni *et al.*, 1979; Zilli, 1996). Being mainly based on features of the habitus and genital morphology (Obraztsov, 1966), the taxonomy of *Syntomis* moths appears to be outdated and confused, evidently flawed by incorrect delimitation of real species boundaries - here we consider *Syntomis* as a genus distinct from *Amata* on the basis of information provided by Schneider *et al.* (1999). In particular, we focused on *S. marjana*, a species which ranges from Provence, Sicily and continental Italy across the Balkan Peninsula, Ukraine and Southern European Russia eastwards to Northern Caucasus (Freina, 2008; Fibiger *et al.*, 2011). Based on their disjunct distribution, and occasionally on some slight morphological differences in size, male genitalia, wing shape and spotting, some populations have occasionally been considered as distinct species, above all *albionica* (Provence), *quercii* (Apennines), *kruegeri* (Sicily), *marjana* (Balkans) and *sheljuzhkoi* (Caucasus) (Verity, 1914; Turati, 1917; Stauder, 1928-1929; Obraztsov, 1966; Dufay, 1970; Igniatev & Zolotuhin, 2005). Preliminary genetic studies showed substantial divergence between two populations of the taxa *kruegeri* (Mt Pellegrino, Palermo, Sicily) and *marjana* (Stari Grad, Croatia), which were then not considered to be conspecific (Cianchi *et al.*, 1980; Bullini *et al.*, 1981). Nevertheless, this genetic evidence has subsequently been challenged by Freina & Witt (1987) and was not considered anymore in recent taxonomic reviews, which recognized only a single widely distributed species with more subspecies (Freina, 2008; Fibiger *et al.*, 2011). Such ongoing discordance on the systematics of the group claims for more comprehensive genetic investigations.

Here we provide the first extensive assessment of the genetic variation across a number of Euro-Anatolian *Syntomis* taxa. Firstly, we assessed the genetic variation at both nuclear and mitochondrial markers within the *S. marjana*-*kruegeri* complex and in another two closely related, sympatrically occurring species, *S. phegea* and *S. ragazzii*; then, we performed ordination-based clustering analyses and used species delimitation methods to circumscribe putative species; finally, we compared the estimated time of divergence among sympatric and allopatric taxa. The aim of this paper is thence to clarify the taxonomy and systematics of a number of Western Palaearctic *Syntomis* taxa and disclose the extent of cryptic diversity within a mimicry complex occurring in temperate areas.

## Materials and Methods

### Sampling and laboratory procedures

The sampling strategy was designed to capture the most of geographic variation from the *Syntomis marjana* -*kruegeri* complex, *S. ragazzii* and *S. phegea*. We analysed a total of 1150 specimens from 34 localities representing most of the described subspecies (we did not analyse *S. m. odessana, S. m. sheljuzhkoi* and *S. p. phegea*). Details about sampling localities and sample sizes are given in Tab. 1 and Fig. 1. Specimens were preliminarily identified following the characters described in Obraztsov (1966), Freina & Witt (1987) and Fibiger *et al.* (2011). Biological tissue was stored at −80° until subsequent analyses. We also obtained two dry specimens from the related species *S. nigricornis* and *S. aequipunta* for use in the phylogenetic analysis (see below).

**Tab. 1.**
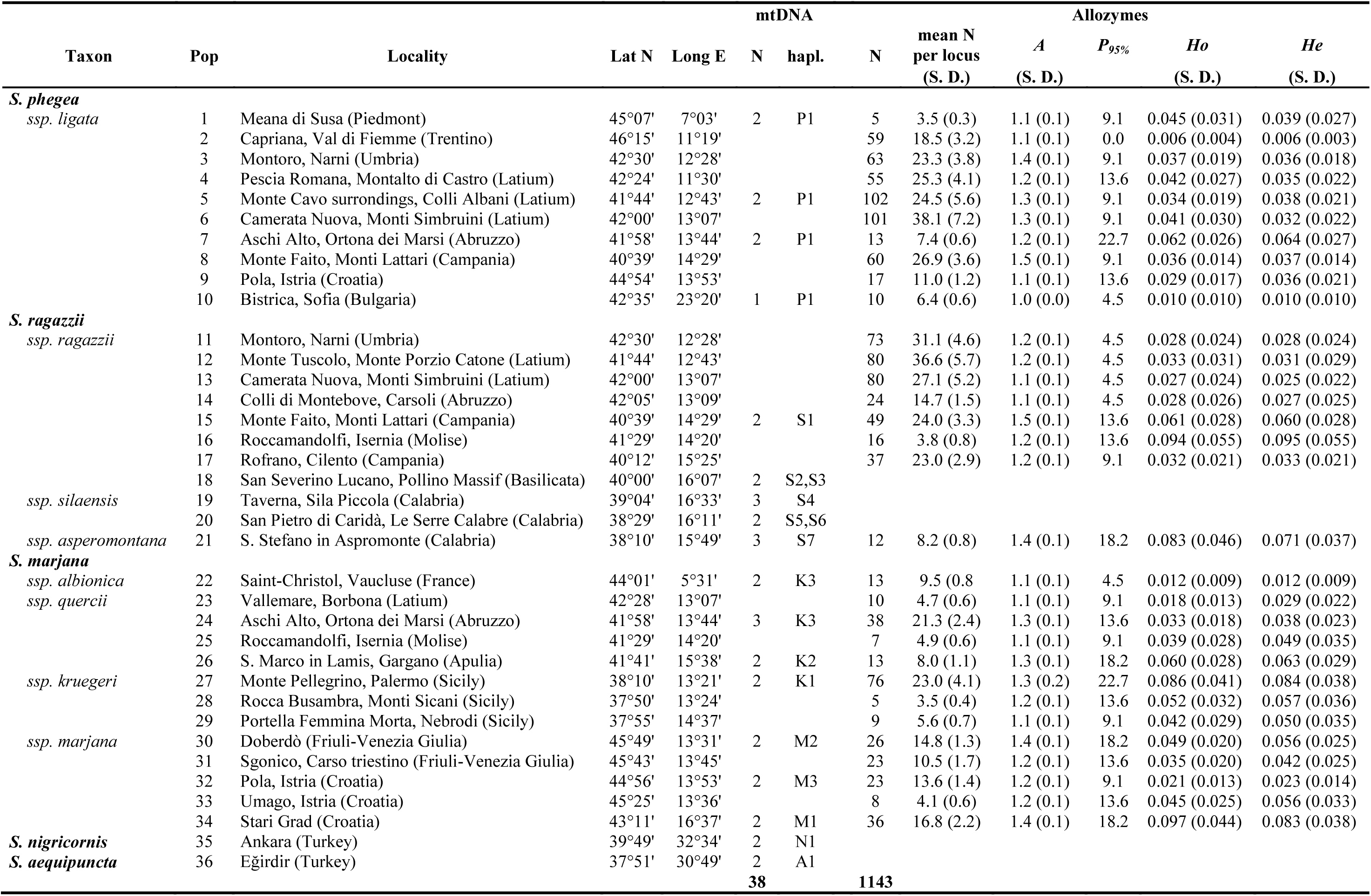
Geographical location of the 36 Western Palearctic *Syntomis* sampling sites, sample size, mtDNA haplotype number and type, estimates of genetic variability at allozyme loci for each population: A, mean number of alleles per locus; P(95%), percentage of polymorphic loci (the most common allele does not exceed 0.95); Ho and He, observed and expected heterozygosity, respectively (with standard deviation). N, sample size.

**Fig. 1.**
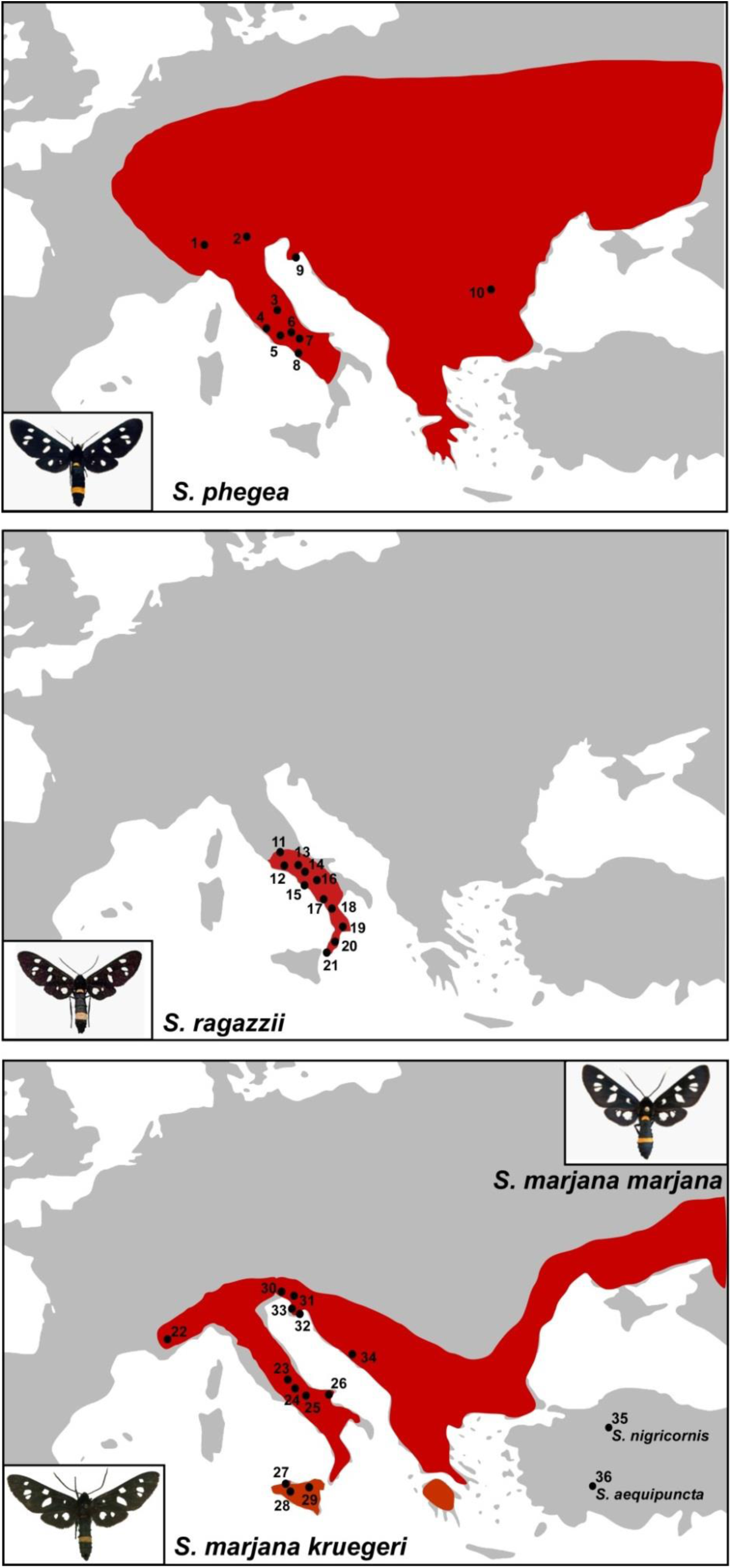
Geographical distribution of *Syntomis phegea*, *S. ragazzii* and *S. marjana sensu lato* in the Western Palearctic region and geographical locations of the 36 populations sampled. Locations are numbered as in Tab. 1. The map was drawn using the software Canvas 11 (ACD Systems of America, Inc.).

Standard horizontal starch gel (10%) electrophoresis was performed to analyse the genetic variation at 22 allozyme loci: α-glycerol phosphate dehydrogenase (*α-Gpdh*), malate dehydrogenase (*Mdh-1* and *Mdh-2*), isocitrate dehydrogenase (*Idh-1* and *Idh-2*), 6-phosphogluconate dehydrogenase (*6Pgdh*), glyceraldehyde-3-phosphate dehydrogenase (*Gapdh*), superoxide dismutase (*Sod-1* and *Sod-3*), xanthine dehydrogenase (*Xdh*), hexokinase (*Hk-1* and *Hk-2*), adenylate kinase (*Adk*), esterase (*Est-1*, *Est-2* and *Est-5*), acid phosphatase (*Acph*), aldolase (*Ald*), aconitase (*Aco*), triosephosphate isomerase (*Tpi*), mannose phosphate isomerase (*Mpi*), and phosphoglucomutase (*Pgm*).

Electrophoretic techniques and staining procedures followed Selander *et al.* (1971), Ayala *et al.* (1972) and Harris & Hopkinson (1976), with minor modifications. Alleles were numbered according to their mobility (expressed in mm) with respect to the most common allele (named 100) in a reference population (*S. phegea* from Camerata Nuova, Central Italy).

DNA was extracted from the legs of a subsample of available specimens (Tab. 1) following the standard cetyltrimethylammonium-bromide (CTAB) protocol (Doyle & Doyle, 1990). A fragment from the mitochondrial Cytochrome Oxidase I gene (*COI*) was amplified and sequenced. The polymerase chain reaction (PCR) primers were REVNANCY (5’-GAA GTT TAT ATT TTA ATT TTA CCG GG-3) and PAT2K837 (5’-TCC ATT ACA TAT AAT CTG CCA TAT TAG-3’) from Simon *et al.* (1994). Amplifications were performed in a 10-μL reaction volume containing MgCl_2_ (2 mM), four dNTPs (0.2 mM each), two primers (0.2 μM each), the enzyme *Taq* polymerase (0.5 U, Promega), its reaction buffer (1X, Promega) and 10–100 ng of DNA template. PCR runs were conducted following an initial step at 95°C for 5 min, then 32 cycles at 94°C for 1 min, 45 sec at 57°C (*COI*) and 1 min at 72°C, followed by a single final step at 72°C for 5 min. Purification and sequencing of PCR products were conducted on both strands by Macrogen Inc. (http://www.macrogen.com), using an ABI PRISM^®^ 3730 sequencing system (Applied Biosystems). All sequences have been deposited in GenBank (accession numbers: XXXX).

### Allozyme data analysis

The effective number of alleles per locus was calculated with software GENALEX (Peakall & Smouse, 2006). Allele frequencies, mean observed and expected heterozygosity and proportion of polymorphic loci were computed for each sampling site (population) using BIOSYS-2 (Swofford & Selander 1999). BIOSYS-2 was also used to evaluate departures from the expected Hardy-Weinberg equilibrium for each locus at each sampling site, and the linkage equilibrium between each pair of loci after application of the Bonferroni correction for multiple tests. As no departures from Hardy-Weinberg equilibria were observed, indicating occurrence of panmixia in all populations, we conducted a population-based assessment of genetic affinities.

A pairwise matrix of unbiased Nei’s genetic distances (Nei, 1978) among all population samples was computed by BIOSYS-2. Clustering of populations was assessed via Principal Coordinate Analysis (PCoA) of Nei’s distances among populations, as implemented in GENALEX. PCoA-defined clusters were then compared with candidate taxa and a matrix of fixed differences (i.e. fully diagnostic loci) among them was generated.

### Sequence data analysis, species delimitation and phylogenetic reconstruction

Electropherograms of sequence data were visually checked using FinchTV 1.4.0 (Geospiza Inc.), and sequences were aligned using Clustal X 2.0 (Larkin *et al.*, 2007). Nucleotide variation was assessed using MEGA 6.0 (Tamura *et al.*, 2013); haplotype and nucleotide diversity were estimated using DnaSP 5.10 (Librado & Rozas, 2009).

We applied three different species delimitation methods which are traditionally employed to delimit groups of sequences that potentially correspond to distinct species: (i) Automatic Barcode Gap Discovery (ABGD; Puillandre *et al.*, 2012a), that detects gaps in the distribution of pairwise genetic distances, assuming that it corresponds to a threshold between intra- and inter-specific distances; (ii) General Mixed Yule Coalescent model (GMYC; Pons *et al.*, 2006), which starts from an ultrametric tree and tests whether the branching rates fit better with a coalescent model or a speciation model, using the transition point between speciation and coalescence to delimit species; and (iii) Bayesian implementation of the Poisson Tree Processes (bPTP; Zhang *et al.*, 2013), which also compares speciation and coalescent models but relies on the substitution rates calculated for each node instead of the branching rates. These three methods were chosen because they proved to be effective in recognizing cryptic species either in large and small groups of sequences and are among the most widely used with mtDNA data, also in single gene analyses (Fujita *et al.*, 2012; Puillandre *et al.*, 2012a, 2012b; Zhang *et al.*, 2013; Schwarzfeld & Sperling, 2015).

ABGD analysis was performed on the ABGD webserver platform (http://www.abi.snv.jussieu.fr/public/abgd/abgdweb.html), using the default parameters (gap width X = 1.5, prior intraspecific divergences from P = 0.001 to P = 0.1, with 20 steps) and the Kimura-2-parameter (K80) model to compute a pairwise genetic distance matrix. The phylogenetic tree used as input for the GMYC and bPTP analyses was generated using the Bayesian method implemented in BEAST 1.8.1 (Bayesian evolutionary analysis by sampling trees; Drummond *et al.*, 2012). We selected the Yule pure-birth speciation model as tree prior, a strict clock model, and the best-fit model of molecular evolution estimated by Jmodeltest 2.1.3 (Darriba *et al.*, 2012) under the Bayesian Information Criterion, i.e. the TIM2 transitional model with a proportion of invariant sites (+I); the Markov chain Monte Carlo (MCMC) length was 10 million generations, sampling trees every 1000 generations. The independence of the effective sample size (ESS values »200) for the estimated parameters was evaluated using Tracer 1.6, after removing the first 10% of samples as burn-in. The consensus tree was then generated by TreeAnnotator 1.8.1 (BEAST package), using the maximum clade credibility criterion, after removing the first 1000 sampled trees as burn-in. We included in the analysis also the sequences of *S. nigricornis* and *S. aequipuncta*. The GMYC analysis was then ran using the *splits* R package (Ezard *et al.*, 2015), applying both the single threshold and the multiple threshold methods. bPTP analysis was conducted on the webserver platform (https://species.h-its.org/ptp/), with a 100000 MCMC length and a 10% burn-in.

The phylogenetic relationships among the candidate species was inferred using maximum likelihood (ML) and Bayesian methods. The ML-tree was generated using the algorithm implemented in IQTREE (Nguyen *et al.*, 2014) applying the TIM2+I model of substitution; the robustness of the inferred tree topology was assessed using the non-parametric bootstrap method with 1000 pseudo-replicates and the SH-like approximate likelihood ratio test (SH-aLRT), also with 1000 bootstrap replicates. The Bayesian phylogenetic analysis was performed using the time-calibrated procedure implemented in BEAST 1.8.1 (Drummond and Rambaut, 2007), in order to get a comparative framework of the times of divergence among the candidate species and to get an approximate historical contextualization of the speciation events. In absence of internal calibration points derived from fossil data or geologic events directly linked to the evolutionary history of *Syntomis*, to time-calibrate the tree we used an uncorrelated relaxed clock model and a fixed substitution rate of 0.0015 per site per million of years, i.e. the mean COI substitution rate estimated for arthropods by Brower *et al.* (1994). Despite the use of non-fossil-based calibrations is usually discouraged (Schenk, 2016), this option is often the only available for most of soft-bodied invertebrates, and continues to be extensively used in dating divergences in insects, including Lepidoptera (e.g. Hinojosa *et al.*, 2018). We chose the substitution rate by Brower (1994) among the several available ones (Papadopoulou *et al.*, 2010) since it falls in the middle of the ranges estimated for this gene fragment in insects, and it is so far the most used, facilitating comparative inferences. The Yule pure-birth speciation model was chosen as tree prior, and the TIM2+I as substitution model. Two independent runs were performed, each with a Markov chain Monte Carlo (MCMC) length of 10 million generations, with sampling every 1000 generations. The independence of the estimated parameters (ESS values >200) and the convergence between runs were evaluated using Tracer 1.6, after removing the first 10% of samples as burn-in. The two runs were combined using LogCombiner 1.8.1 (BEAST package), and an annotated maximum clade credibility tree was computed with TreeAnnotator 1.8.1.

## Results

### Allozyme data analysis

Among the 22 loci analysed, four (*Sod-1*, *Sod-3*, *α-Gpdh*, *Gapdh*) resulted monomorphic in all studied populations. The allele frequencies at the 18 polymorphic loci are given in the Supplementary Table 1. The overall number of alleles observed at each polymorphic locus varied from two to nine (*Pgm*). No departures from both the Hardy-Weinberg and linkage equilibria were observed (no significant value after the Bonferroni correction). The mean sample size per locus, the effective number of alleles and the observed and expected heterozygosity for each population are shown in Tab. 1. The mean sample size ranged from 3.5 (pop. 28) to 38.1 (pop. 6), the average number of alleles per locus varied from 1.1 (several samples) to 1.5 (pop. 8, *S. phegea* from Monte Faito, and pop. 15, *S. ragazzii* from Monte Faito), the mean observed heterozygosity ranged from 0.006 (pop. 2, *S. phegea* from Capriana) to 0.097 (pop. 34, *S. marjana* from Stari Grad), the mean expected heterozygosity ranged from 0.006 (pop. 2, *S. phegea* from Capriana) to 0.095 (pop. 16, *S. ragazzii* from Roccamandolfi).

The pairwise matrix of population genetic distances D (Nei, 1978) is given in Supplementary Tab. 2, whereas a heatmap of the D values is shown in Fig. 2. The inspection of the heatmap revealed four main clusters of populations with lower values of genetic distance, which coincided with populations of *S. phegea*, *S. ragazzii*, *S. m. marjana*, and *S. m. kruegeri* + *S. m. quercii* + *S. m. albionica*. The mean genetic distances observed within these groups are 0.002, 0.013, 0.013 and 0.017, respectively, which are values commonly observed at the intraspecific level (Bullini & Sbordoni, 1980). The mean genetic distances among these groups range from 0.338 (*S. phegea* vs. *S. m. marjana*) to 0.629 (*S. ragazzii* vs. *S. m. kruegeri* + *S. m. quercii* + *S. m. albionica*); the genetic distance between *S. m. marjana* and *S. m. kruegeri* + *S. m. quercii* + *S. m*. *albionica* was 0.432; these values agree with those commonly observed among closely related species (Bullini & Sbordoni, 1980). The highest values of genetic distance observed within groups are 0.076, between southern (pop. 34, Stari Grad) and northern (pop. 31, Sgonico) population of *S. m. marjana*, 0.075, between *S. m. kruegeri* (pop. 28, Rocca Busambra) and *S. m. quercii* (pop. 24, Aschi Alto), and 0.063, between *S. r. asperomontana* (pop. 21, S. Stefano in Aspromonte) and *S. r. ragazzii* (pop. 16, Roccamandolfi); these values agree with those commonly observed among subspecies (Bullini & Sbordoni, 1980). A summarized matrix of the mean genetic distances observed within and between these groups is given in Tab. 3.

**Tab. 2.** Results from the species delimitation analyses using the ABGD, GMYC and bPTP methods on mtDNA haplotypes (details in the text). Site and haplotype codes follow Tab. 1. Different species are coded with different letters (A-F).

| Taxon | Site | haplotypes | ABGD |  | GMYC |  | PTP |
| --- | --- | --- | --- | --- | --- | --- | --- |
|  |  |  | primary partition | secondary partition | single threshold | multiple threshold |  |
| *S. phegea ligata* | 1 | P1 | A | A | A | A | A |
| *S. ragazzi ragazzii* | 15 | S1 | B | B | B | B | B |
| *S. ragazzi ragazzii* | 18 | S2, S3 | B | B | B | B | B |
| *S. ragazzi silaensis* | 19 | S4 | B | B | B | B | B |
| *S. ragazzi silaensis* | 20 | S5, S6 | B | B | B | B | B |
| *S. ragazzi asperomontana* | 21 | S7 | B | B | B | B | B |
| *S. majana albionica* | 22 | K3 | C | C1 | C | C1 | C1* |
| *S. majana quercii* | 24 | K3 | C | C1 | C | C1 | C1* |
| *S. majana quercii* | 26 | K2 | C | C1 | C | C1 | C1* |
| *S. majana kruegeri* | 27 | K1 | C | C2 | C | C2 | C2* |
| *S. majana marjana* | 30 | M2 | D | D1 | D | D1 | D1 |
| *S. majana marjana* | 32 | M3 | D | D1 | D | D1 | D1 |
| *S. majana marjana* | 34 | M1 | D | D2 | D | D2 | D2 |
| *S. nigricornis* | 35 | N1 | E | E | E | E | E |
| *S. aequipuncta* | 36 | A1 | F | F | F | F | F |

\* not supported

**Tab. 3.** Reduced matrix of the pairwise mean genetic distance (D Nei 1978) between (below the diagonal) and within (on the diagonal) the four genetic clusters identified by allozyme loci, and number of 100% diagnostic loci between them (above the diagonal).

|  | *S. phegea* | *S. ragazzii* | *S. m. kruegeri* +<br>*S. m. quercii* +<br>*S. m. albionica* | *S. m. marjana* |
| --- | --- | --- | --- | --- |
| *S. phegea ligata* | 0.002<br>(0.000-0.008) | 7 | 9 | 4 |
| *S. ragazzii ragazzi* + *S. r. asperomontana* | 0.536<br>(0.455-0.591) | 0.013<br>(0.000-0.034) | 7 | 5 |
| *S. m. kruegeri* + *S. m. quercii* + *S. m. albionica* | 0.629<br>(0.557-0.706) | 0.597<br>(0.516-0.647) | 0.017<br>(0.002-0.048) | 5 |
| *S. m. marjana* | 0.338<br>(0.290-0.392) | 0.523<br>(0.429-0.629) | 0.432<br>(0.345-0.518) | 0.013<br>(0.003-0.031) |

**Fig. 2.**
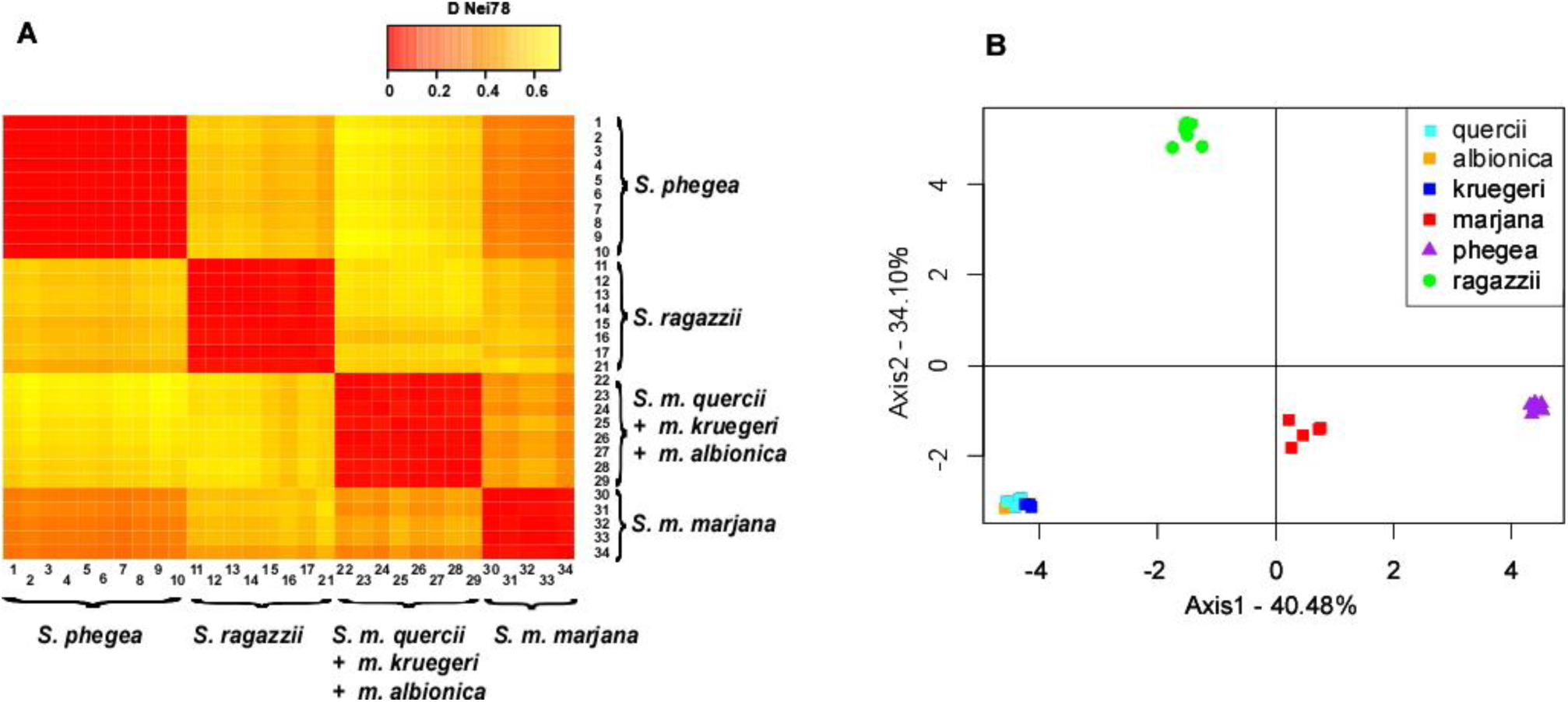
A) Heatmap representing the pairwise matrix of population genetic distance (D Nei 78) among the Western Paleartic *Syntomis* populations analysed in this study, based on 22 allozyme loci; warmer colours indicate higher genetic identity. B) Principal coordinate analysis of the 31 analysed populations, based on the unbiased Nei genetic distance (D Nei 78), calculated on 22 allozyme loci; number in parentheses refer to the proportion of variance explained by the first two principal coordinates. The graph was drawn using the software Canvas 11 (ACD Systems of America, Inc.).

The ordination-based clustering analysis resulting from PCoA also defined four main clusters of populations (Fig. 2). The first and second PCoA axes explained 40.48% and 34.10% of total variance, respectively. Populations of *S. m. kruegeri* + *S. m. quercii* + *S. m. albionica* were grouped together in a distinct and well-defined cluster with respect to populations of *S. m. marjana*, which were also combined into a single cluster; the other two clusters coincided with populations of *S. phegea* and S*. ragazzii*.

The four clusters were also defined by some fully diagnostic loci, resumed in Tab. 3. We found four 100% diagnostic loci between *S. phegea* and *S. m. marjana*, five between *S. marjana* and *S. m. kruegeri* + *S. m. quercii* + *S. m. albionica*, five between *S. m. marjana* and *S. ragazzii*, seven between *S. ragazzii* and *S. m. kruegeri* + *S. m. quercii* + *S. m. albionica*, seven between *S. ragazzii* and *S. phegea*, and nine between *S. phegea* and *S. m. kruegeri* + *S. m. quercii* + *S. m. albionica*.

### Sequence data

We successfully amplified and sequenced a 781 bp fragment from the final section of the mitochondrial COI gene from 38 *Syntomis* individuals. No indels, stop codons or non-sense codons were observed. We found 16 different haplotypes defined by 81 (10.3%) variable positions, of which 68 (8.7%) were parsimony informative. The mean haplotype diversity (h) and nucleotide diversity (π) values for this dataset were 0.950 (±0.021 SD) and 0.032 (±0.008 SD), respectively. A full list of the haplotype founds within each studied population, with the respective GenBank accession numbers, is presented in Tab. 1.

The results from the species delimitation analyses using ABGD, GMYC and bPTP are resumed in Tab. 2. ABGD recognized a gap in the pairwise distance distribution between 0.013 and 0.035. Taking the prior maximal distance (P_max_) threshold within this gap, ABGD computed a primary partition with six groups, coinciding with *S. phegea*, *S. ragazzii*, *S m. marjana*, *S. m. kruegeri* + *S. m. quercii* + *S. m. albionica*, *S. nigricornis* and *S. aequipuncta*. The secondary partition, achieved by the recursive analysis, recognized the same groupings when using P_max_ values included in the gap; further splits were recognized for lower values: northern *S. m. marjana* from southern *S. m. marjana* (for P_max_ = 0.0077), and *S. m. kruegeri* from *S. m. albionica* + *quercii* (for P_max_ = 0.0046).

GMYC single-threshold analysis rejected the null hypothesis of a coalescent model (likelihood ratio 6.415; LR test: 0.04, significant) and recognized six candidate species, which coincided with those recovered from ABGD analyses. The multiple-threshold analysis also rejected the coalescent model (likelihood ratio 6.873; LR test: 0.03, significant), but suggested further splits between northern and southern *S. m. marjana* and between *S. m. kruegeri* and *S. m. albionica* + *quercii* (as in ABGD analysis).

Finally, bPTP returned eight putative species, coinciding with *S. nigricornis* (support = 1.0), *S. aequipuncta* (support = 1.0), *S. phegea* (support = 1.0), *S. ragazzii* (support = 0.79), northern *S. m. marjana* (support = 0.87), southern *S. m. marjana* (support = 0.94), *S. m. kruegeri* (not supported) and *S. m. albionica* + *quercii* (not supported).

The ML-tree retrieved by IQTREE (log-likelihood score: −1608.30, s.e. 61.93) recognized six main branches, which were consistent with results obtained from the species delimitation methods (Fig. 3A). The sequences of the taxa *kruegeri*, *quercii* and *albionica* clustered together (high support = 96.4/98%), and resulted the sister clade (high support = 87.4/88%) of *S. m. marjana* clade (high support = 98/100%). The sequences of *S. ragazzii* form a unique supported cluster (high support = 96.5/79%), which resulted more related to *S. nigricornis* (moderate support = 80/66%) than *S. phegea*, whereas *S. phegea* resulted more related to *S. aequipuncta* albeit with weak support (77/54%).

**Fig. 3.**
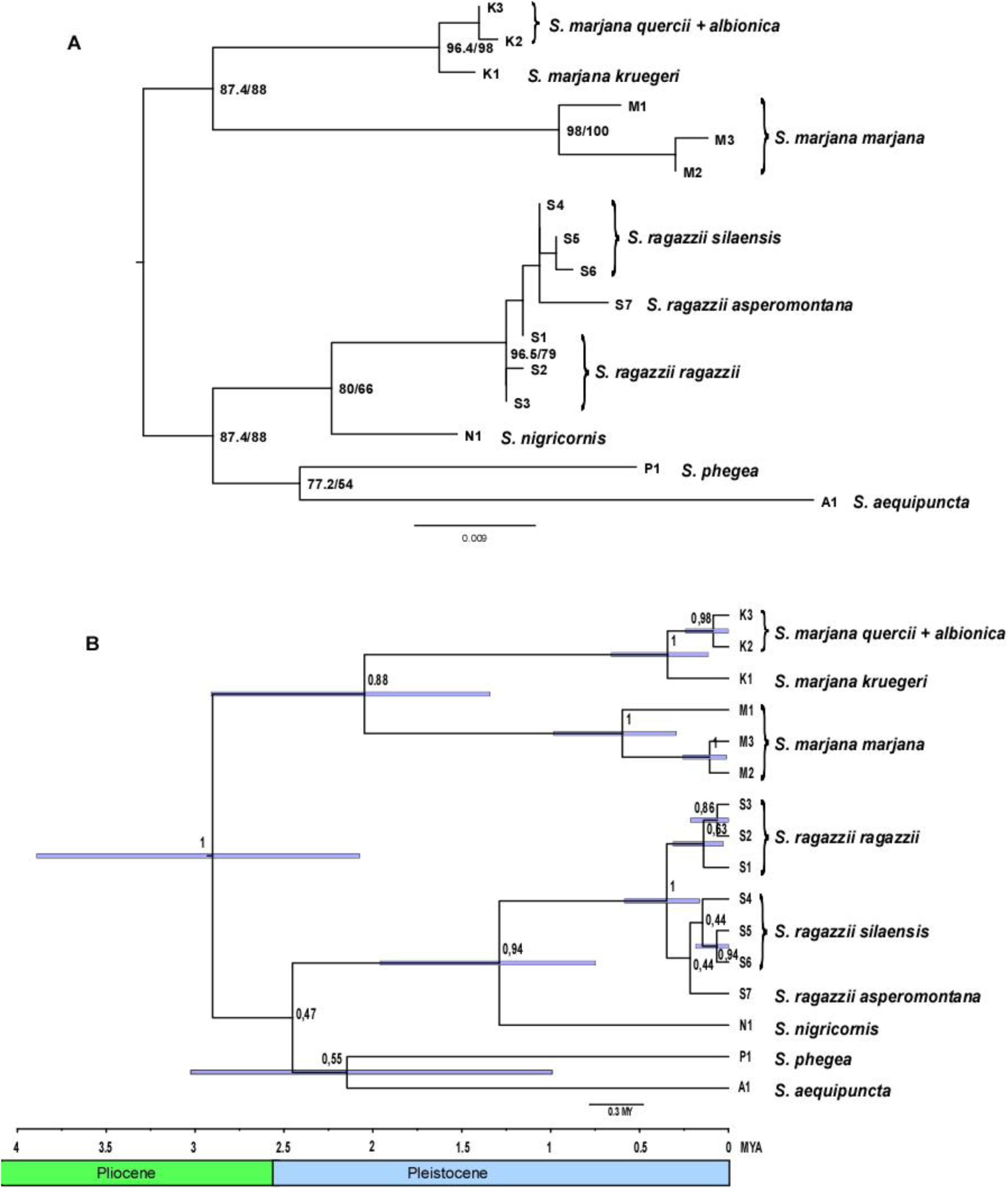
Phylogenetic relationships among the 16 mtDNA haplotypes observed. (A) Maximum likelihood tree retrieved by the analysis in IQTREE; support values at the relevant nodes are SH-aLRT support (%) and standard bootstrap support (%) based on 1000 replicates. (B) Maximum clade credibility tree recovered by the Bayesian analysis in BEAST, showing the divergence time from the most recent common ancestor (TMRCA) for the major clades; node bars (in blue) represent 95% highest posterior density (HPD) intervals for node ages; posterior probabilities for each node are also shown; MYA: million years ago. The graph was drawn using the software Canvas 11 (ACD Systems of America, Inc.).

The time-calibrated phylogenetic analysis conducted in BEAST returned two congruent runs that fully converged to a stationary distribution with satisfactory ESS values (>>200) for all the parameters of interest. A chronogram based on the maximum clade credibility (MCC) tree is presented in Fig. 3B. The tree topology was consistent with both the ML-tree and the species delimitation analyses, as it showed six main branches coinciding with *S. phegea*, *S. aequipuncta*, *S. nigricornis*, *S. ragazzii*, *S. m. kruegeri* + *S. m. quercii* + *S. m. albionica* and *S. m. marjana*. The time to the most recent common ancestor (MRCA) for all the analysed taxa was estimated at 2.87 million years ago (mya) (95% HPD:1.87 – 4.12); that between *S. m. kruegeri* + *S. m. quercii* + *S. m. albionica* and *S. m. marjana* at 1.99 mya (95% HPD: 1.11 – 2.96), roughly the same as that between *S. phegea* and *S. aequipuncta* (mean: 1.92 mya, 95% HPD:0.93 – 3.05), and higher than that estimated for *S. ragazzii* and *S. nigricornis* (mean: 1.29, 95% HPD: 0.66 – 2.07). The estimated TMRCA for the clades coinciding with *S. m. marjana*, *S. m. kruegeri* + *S. m. quercii* + *S. m. albionica* and *S. ragazzii* were 0.61 mya (95% HPD: 0.26 – 1.1), 0.37 mya (95% HPD: 0.10 – 0.76) and 0.36 mya (95% HPD: 0.08 – 0.67), respectively.

## Discussion

Both mitochondrial and nuclear markers consistently supported the existence of two distinct, highly differentiated, genetic clusters within *S. marjana*, distributed West and East of the Adriatic Sea. The extent of the genetic differentiation between these two clusters and the estimated time of their divergence are of the same magnitude of those occurring among other well-recognized species of this complex. Furthermore, all species delimitation methods clearly point to these two genetic clusters as distinct species. These results are consistent with preliminary evidence showing substantial genetic divergence between the taxa *kruegeri* and *marjana* s. str. (Cianchi *et al.*, 1980, Bullini *et al.*, 1981), and suggest to consider the populations of the lineage *kruegeri* + *quercii* + *albionica* as a distinct species, under the oldest available name taking priority, that is *S. quercii* Verity, 1914 (see below). Our data also highlight further genetic sub-structuring at intraspecific level, suggesting a role for recent Pleistocene events in triggering population-level genetic differentiation.

The taxa *kruegeri*, *marjana* and *quercii* were originally described as aberrations or subspecies of other species, the first two of *S. phegea* (Ragusa, 1904; Stauder, 1913), the last one of *S. mestralii*, a species from the Middle East. Subsequently, *kruegeri* and *marjana* were both considered to be valid species by Turati (1917), who however ranked *quercii* as a subspecies of *marjana*. However, the strong similarity of these species in habitus, habitat preferences (dry grasslands) and phenology, together with the difficulties in identifying fully reliable diagnostic characters, led most authors to combine them into a unique polytypic species (Obraztsov, 1966; Bertaccini *et al.*, 1997; Igniatev & Zolotuhin, 2005; Freina, 2008; Fibiger *et al.*, 2011). Despite their strong similarity, we detected substantial genetic differentiation between *S. marjana* and *S. quercii* (as aforementioned comprising herein also the taxa *albionica* and *kruegeri*) that can be attributed to a speciation event occurred at some point in the Early Pleistocene or before. Our estimate of the divergence time between these two lineages is probably biased by the use of a non-fossil-based calibration of the molecular clock (Papadopoulou *et al.*, 2010; Schenk, 2016), but it provides an approximate timing of their split. Indeed, the strong changes in bioclimatic conditions that characterized the Plio-Pleistocene transition prompted speciation in several temperate species inhabiting the Mediterranean peninsulas (Hewitt, 1996, 2011), including Lepidoptera (Schmitt, 2007), and likely affected also the evolutionary history of the European *Syntomis*. These changes consisted of a generalized increase of aridity and a reduction of the forest habitats in Southern Europe (Hewitt, 2011), followed by the beginning of glacial cycles, that resulted in alternate expansion and retreat of xeric and forest habitats (Faquette *et al.*, 1999; Suc & Popescu, 2005; Abrantes *et al.*, 2010). Most European *Syntomis* are thermophilous (with the partial exception of *S. phegea*), occur in dry habitats, and are distributed in South-Eastern Europe with a number of relatives in steppe areas from the Middle-East to Central Asia, that likely represents the centre of origin of this group (“*phegea*-Gruppe” of Obraztsov, 1966). The common ancestor of *S. marjana* and *S. quercii* might have benefited from the increase of aridity in Western Europe at the end of Pliocene (Malatesta, 1985; Faquette *et al.*, 1999) by spreading over South-Eastern Europe. Then, with the advent of the first glacial cycle at the beginning of Pleistocene, the cooler conditions and the spreading of broad-leaved forests in the Italian and Balkan Peninsulas (Abrantes *et al.*, 2010) could have trapped the ancestor in the south of both peninsulas, where populations would have found relatively hot and dry refugia. This vicariance process has been claimed as the most likely explanation for the existence of several sibling species pairs with allo-parapatric distribution in the Italian and Balkan peninsulas (Racheli & Zilli, 1985). The Isonzo river in NE Italy and Slovenia, which virtually separates the distribution of these species, has been considered as a suture zone and biogeographical boundary for several vertebrate and invertebrate species (Taberlet *et al.*, 1998; Hewitt, 1999). However, in our case is hard to define a boundary between these two species because most of the historically known populations just west of that river have not been found anymore over the last 50 years and could not be genotyped. Further genetic assays on museum specimens are therefore required.

Pleistocene climatic changes do also support the origin of genetic sub-structuring of conspecific populations. Despite our data do not allow fine-scale phylogeographic inferences, they clearly reveal genetic sub-structuring in *S. quercii*, *S. ragazzii* and *S. marjana*, which can be explained by the interaction between climate fluctuations and the physiographic heterogeneity of the Italian and Balkan peninsulas. We found genetic differentiation between the Sicilian and the Apennine populations of *S. quercii*, between the Central and southernmost Apennine populations of *S. ragazzii*, and between the northern and the southern populations of *S. marjana* s. str. (Fig. 3 and Supplementary Tab. 2. Substantial intraspecific differentiation between Sicilian and mainland populations and between the southernmost and Central Apennine populations is commonly observed in phylogeographic studies of numerous Italian species (Canestrelli *et al.*, 2008,; 2010; Chiocchio *et al.*, 2019; Scalercio *et al.*, 2019). These patterns are ascribable to the presence of several geographic discontinuities that acted as ecological and physiographic barriers at some point in the past, after a “refugia-within-refugia” scenario (Gomez & Lunt, 2007). During Pleistocene, the southernmost section of the Italian Peninsula and Sicily have been affected by several glacio-eustatic oscillations of the sea level that deeply influenced the genetic structure of populations (Bonfiglio *et al.*, 2002). This scenario also fits well with the genetic structure observed in *S. ragazzii* and *S. quercii*. In contrast, no apparent paleogeographic evidence supports the genetic differentiation observed within *S. marjana* in a relatively restricted area along the NE Adriatic with no evident orographic discontinuity. However, this area is interested by geological discontinuities (Cvetkovic *et al.*, 2015) that affect ground water availability and, as a consequence, vegetation and bioclimatic conditions (DMEER, 2017). These bioclimatic discontinuities likely represented ecological barriers for *S. marjana* populations which may have limited their gene flow, at least during the last part of the Pleistocene.

In the phylogenetic relationships outlined by mitochondrial DNA, noteworthy is the recovery of *S. ragazzii* as more closely related to the Anatolian-Caucasian *S. nigricornis* than *S. phegea*, despite *S. ragazzii* and *S. phegea* were long considered as sibling species and can still occasionally hybridize (Sbordoni *et al.*, 1982). Vicariance between Southern Apennine and Anatolian taxa is fairly common in Lepidoptera and can be ascribed to several biogeographic processes (Racheli & Zilli, 1985) other than a direct geological connection between the two regions, the last of which is estimated in the late Miocene (Rögl *et al.*, 1999). Interestingly, the divergence between these two species has been estimated to be more recent than those between the other sister pairs. However, more samples of *S. nigricornis* and other middle-eastern taxa are strongly needed to infer the whole biogeographic history of this connection. Furthermore, the weak support values recovered for the relationship of *S. phegea* and *S. aequipuncta* with the lineages of *quercii*-*marjana* and *ragazzii*-*nigricornis* stress the need to broaden the analysis to more genetic markers and species in order to fully resolve the evolutionary history of the Western Palearctic *Syntomis* (see also Przybyłowicz *et al*., 2019).

Our results highlight that in Euro-Anatolian *Syntomis* the degree of genetic divergence is not related with that of phenotypic divergence, as expected for species involved into mimicry complexes. These moths happen thus to be at the crossroad between independent evolutionary divergence and constraints limiting their phenotypic diversification (Leimar *et al.*, 2012). Indeed, species belonging to the same Müllerian mimicry ring share the costs of predation but they have to be perceived very similar by predators. As a consequence, the external appearance of these species is expected to be under strong ecological selection to maintain a common signal. Accordingly, different set of characters (e.g. morphological, genetic, physiological, behavioural, etc.) will reveal their interplay with the various evolutionary forces, that is in turn affected by the allopatric/sympatric occurrence of populations of these moths with those of other co-mimics. Many *Syntomis* species are sympatric and coexist in the same or in really close biotopes, at least to the scale of bird predators’ home ranges. *Syntomis quercii* and *S. marjana* have allo-parapatric distribution, but they both overlap with *S. phegea* (except in Sicily, where the latter is absent, all records hitherto being misidentifications), which is extremely abundant and has already shown to be a Batesian model also for some distantly related co-mimics, like the yellow ephialtoid form of *Zygaena ephialtes* (Sbordoni *et al.*, 1979). In this context, departure of pattern from a successful aposematic signal would be strongly counter-selected during and after speciation. Strong ecological constraints would then be responsible for morphological stasis, in the face of substantial genetic divergence, thus eventually leading to the evolution of so-called cryptic diversity (Bickford *et al.*, 2007; Struck *et al.*, 2018). Interestingly, such stasis extends also to structural features of the genitalia, all fairly homogeneous across the group, which cannot evidently be modelled by predation or other external agents. This circumstance, which sharply contrasts with the striking genital diversification seen in other groups of externally almost indistinguishable moth species (e.g. the sympatric species pairs *Grammodes geometrica-G. occulta*, *Dysgonia algira-D. torrida*, *Noctua fimbriata-N. tirrenica* and *Cilix glaucata-C. hispanica*, to name just a few), suggests that genitalia in *Syntomis* maintain stable configurations due of phylogenetic inertia even after speciation, as they do not play any species isolation role.

As a matter of fact, mechanical compatibility of genitalia between different *Syntomis* species has been proved by the detection of occasional hybrids *S. phegea x S. ragazzii* (Sbordoni *et al.*, 1982) and heterospecific fertile matings *S. phegea x S. marjana* reared from the wild (Rupinpiccolo, NE Italy, A. Zilli, unpublished). Mate recognition and species integrity in *Syntomis* has therefore to rely on other systems than mechanical compatibility but, as seen above, visual stimuli are unfit candidates for that as any departure from the shared pattern would weaken efficacy of their common aposematic signalling. Bickford *et al.* (2007) recognise that cryptic diversity is particularly common in taxa not using visual mating signals. Sheer evidence that *Syntomis* species are unlikely to use visual mate recognition systems is again based on field observations of (pseudo)copulae with completely unrelated and differently patterned moths of a different family, members of the genus *Zygaena* which are part of an alternative mimicry ring based on red and black colours. Such mismatings are not uncommon during peaks of abundance of these moths, when they cluster together on flower heads and get evidently confused by congestion of pheromonal plumes all around and tactile abdominal stimuli which lead them to clasp almost anything within their reach (Lees & Zilli, in press). Breeding experiments run by us (unpublished) are fully compatible with chemically mediated interactions between sexes in *Syntomis* moths, with females typically resting in a calling posture and attracting males from distance. Furthermore, Bendib & Minet (1998) demonstrated presence of female pheromone glands in *S. phegea*, whereas Schneider at al. (1999) have demonstrated that males of all *Syntomis* species bear androconial organs at the base of forelegs. Thus, all evidence points to chemical communication as the main driver of mate recognition in *Syntomis* moths.

### Taxonomic remarks and Conclusions

Our research confirms that in one of the taxonomically most controversial groups of European moths, the Euro-Anatolian *Syntomis*, additional cryptic species are involved. This is the case of a group of populations living in the Italian peninsula, Sicily and Provence that have been noted by different authors as *quercii, kruegeri* and *albionica*, respectively, and will have to be overall combined under the last name, i.e. *Syntomis quercii* Verity, 1914 (**bona sp.**, **stat. nov.**). In fact, when first named by Ragusa (1904), the oldest name *kruegeri* was infrasubspecific and before that it was made available for nomenclature by Turati (1917), Verity’s (1914) description intervened.

Further genetic sub-structuring has been detected in *S. quercii*, *S. ragazzii* and *S. marjana s.s.* Even if it is a common mistake to assign to populations genetically differentiated from nominotypical ones a subspecific rank, as the subspecies concept was not intended to be also evolutionary (see Zilli, 1996, and references therein), it is worth noting that at least as to *S. ragazzii*, its diversification in the southernmost part of its range (Calabrian Apennine) had already been noted with the description of two traditional subspecies (*silaensis* Obraztsov, 1966 and *asperomontana* [Stauder & Turati], 1917). Partially distinct genetic make-ups also characterise the Sicilian populations of *S. quercii*, traditionally recognized as ssp. *kruegeri*.

Research on the Euro-Anatolian taxa of *Syntomis* highlights how the Palaearctic species of this group provide an interesting example to study trade-offs between morphological stasis and genetic divergence in complexes involved into mimicry relationships of Müllerian type. New studies will have to be focused on understanding the genomic basis of speciation and origin of cryptic diversity within this group of moths.

## Supporting information

Allele frequencies at the 18 polymorphic allozyme loci studied

Matrix of genetic distance (Nei, 1978) among the studied population samples

## Supporting information

Additional supporting information may be found online in the Supporting Information section at the end of the article.

**Supplementary table 1**. Allele frequencies at the 18 polymorphic allozyme loci studied.

**Supplementary table 2.** Matrix of genetic distance (Nei, 1978) among the studied population samples.

## Acknowledgments

We would like to thank Mauro Zampiglia for his help on fieldwork. AC also thanks J. Rota for her helpful suggestions. The authors declare that there are no conflicts of interest.

## References

Ayala, F. J., & Powell, J. R. (1972). Allozymes as diagnostic characters of sibling species of *Drosophila*. Proceedings of the National Academy of Sciences, 69, 1094–1096.

Ayala F.J., Powell J.R., Tracey M.L., Mourão C.A. & Pérez-Salas, S. (1972). Enzyme variability in the *Drosophila willistoni* group. IV. Genic variation in natural populations of *Drosophila willistoni*. Genetics, 70, 113–139.

Abrantes, F., Voelker, A., Sierro Sanchez, F., Naughton, F., Rodrigues, T., Cacho, I., Ariztegui, D., Brayshaw, D., Sicre, M-A. & Batista, L. (2010) Paleoclimate variability in the Mediterranean region. The Climate of the Mediterranean Region (ed. by Lionello, P.) pp. 1–86, Elsevier, London.

Bálint, M., Domisch, S., Engelhardt, C.H.M., Haase, P., Lehrian, S., Sauer, J., Theissinger, K., Pauls, S.U. & Nowak, C. (2011) Cryptic biodiversity loss linked to global climate change. Nature Climate Change, 1, 313–318.

Beheregaray, L.B. & Caccone, A. (2007) Cryptic biodiversity in a changing world. Journal of Biology, 6, 9.

Bendib, A. & Minet, J. (1998) Female pheromone glands in Arctiidae (Lepidoptera). Evolution and phylogenetic significance. Comptes Rendus de l’Académie des Sciences, (III) (Sciences de la Vie), 321, 1007–1014.

Bertaccini, E., Fiumi, G. & Provera, P. (1997) Bombici e Sfingi d‟Italia (Lepidoptera: Heterocera). Volume 2. Natura – Giuliano Russo Editore, Bologna.

Bickford, D., Lohman, D.J., Sodhi, N.S., Ng, P.K., Meier, R., Winker, K., Ingram, K.K. & Das, I. (2007) Cryptic species as a window on diversity and conservation. Trends in Ecology & Evolution, 22, 148–155.

Bonfiglio, L., Mangano, G., Marra, A.C., Masini, F., Pavia, M. & Petruso, D. (2002) Pleistocene Calabrian and Sicilian bioprovinces. Geobios, 35, 29–39.

Boppré, M., Vane-Wright, R.I. & Wickler, W. (2017) A hypothesis to explain accuracy of wasp resemblances. Ecology and Evolution, 7, 73–81.

Brower, A.V. (1994) Rapid morphological radiation and convergence among races of the butterfly Heliconius erato inferred from patterns of mitochondrial DNA evolution. Proceedings of the National Academy of Sciences, 91, 6491–6495.

Bullini, L. & Sbordoni, V. (1980) Electrophoretic studies of gene-enzyme systems: microevolutionary processes and phylogenetic inference. Bolletino di Zoologia, 47(suppl.), 95–112.

Bullini, L., Sbordoni, V. & Ragazzini, P. (1969) Mimetismo mülleriano in popolazioni italiane di *Zygaena ephialtes* (L.) (Lepidoptera, Zygaenidae). Archivio Zoologico Italiano, 54, 181–214.

Bullini, L., Cianchi, R., Stefani, C. & Sbordoni, V. (1981) Biochemical taxonomy of the *Amata phegea* complex (Ctenuchidae, Syntominae). Nota lepidopterologica 4, 125–127.

Canestrelli, D. & Nascetti, G. (2008) Phylogeography of the pool frog *Rana (Pelophylax) lessonae* in the Italian peninsula and Sicily: multiple refugia, glacial expansions and nuclear–mitochondrial discordance. Journal of Biogeography, 35, 1923–1936.

Canestrelli, D., Aloise, G., Cecchetti, S. & Nascetti, G. (2010) Birth of a hotspot of intraspecific genetic diversity: notes from the underground. Molecular Ecology, 19, 5432–5451.

Carstens, B.C., Pelletier, T.A., Reid, N.M. & Satler, J.D. (2013) How to fail at species delimitation. Molecular ecology, 22, 4369–4383.

Chiocchio, A., Colangelo, P., Aloise, G., Amori, G., Bertolino, S., Bisconti, R., Castiglia, R. & Canestrelli, D. (2019) Population genetic structure of the bank vole Myodes glareolus within its glacial refugium in peninsular Italy. Journal of Zoological Systematics and Evolutionary Research, (early view).

Cianchi, R., Stefani, C., Sbordoni, V. & Bullini, L. (1980) Ricerche elettroforetiche sulle specie italiane del complesso *Amata phegea* (Lepidoptera, Ctenuchidae): aspetti genetici, tassonomici ed evolutivi. Atti XII Congresso nazionale italiano di Entomologia, 2, 239–242.

Cvetkovic, V., Prelević, D. & Schmid, S. (2016) Geology of South-Eastern Europe. In: Mineral and thermal waters of Southeastern Europe (pp. 1–29). Springer, Cham.

Darriba, D., Taboada, G.L., Doallo, R. & Posada, D. (2012) jModelTest 2: more models, new heuristics and parallel computing. Nature methods, 9, 772.

Doyle, J.J. & Doyle, J.L. (1990) Isolation of plant DNA from fresh tissue. Focus, 12, 39–40.

DMEER (2017). Digital Map of European Ecological Regions. Update 01 Nov 2017. European Environment Agency. http://www.eea.europa.eu/data-and-maps/data/digital-map-of-european-ecological-regions

Drummond, A.J. & Rambaut, A. (2007) BEAST: Bayesian evolutionary analysis by sampling trees. BMC evolutionary biology, 7, 214.

Drummond, A.J., Suchard, M.A, Xie, D. & Rambaut, A. (2012) Bayesian phylogenetics with BEAUti and the BEAST 1.7. Molecular biology and evolution, 29, 1969–1973.

Dufay, C. (1970) *Amata albionica* Dufay, bona species et son ethologie (Lep. Ctenuchidae). Entomops, 17, 31–40.

Ezard, T., Fujisawa, T. & Barraclough, T.G. (2015) SPLITS: SPecies’ LImits by Threshold Statistics. R package version 1.0-18/r45.

Fauquette, S., Suc, J.P., Guiot, J., Diniz, F., Feddi, N., Zheng, Z., Bessais, E. & Drivaliari, A. (1999) Climate and biomes in the West Mediterranean area during the Pliocene. Palaeogeography, Palaeoclimatology, Palaeoecology, 152, 15–36.

Fennessy, J., Bidon, T., Reuss, F., Kumar, V., Elkan, P., Nilsson, M.A., Vamberger, M., Fritz, U. & Janke, A. (2016) Multi-locus analyses reveal four giraffe species instead of one. Current Biology, 26, 2543–2549.

Fibiger, M., Laszló, G.M., Ronkay, G., Ronkay, L., Speidel, W., Varga, Z., Wahlberg, N., Witt, T.J., Yela, J.L., Zahiri, R. & Zilli, A. (2011) Lymantriinae and Arctiinae, including phylogeny and check list of the quadrifid Noctuoidea of Europe. Noctuidae Europaeae 13. Entomological Press, Sorø.

Freina, J.J. de (2008) Über die Biologie, Morphologie, Phänologie und Taxonomie von *Amata (Syntomis) kruegeri* (Ragusa, 1904) (Lepidoptera: Arctiidae, Syntominae, Syntomini). Nachrichten des Entomologischen Vereins Apollo, (N.F.), 28, 97–107.

Freina, J.J., de & Witt, T.J. (1987) Die Bombyces und Sphinges der Westpalaearktis. Forschung & Wissenschaft, München.

Fujita, M.K., Leaché, A.D., Burbrink, F.T., McGuire, J.A. & Moritz, C. (2012) Coalescent-based species delimitation in an integrative taxonomy. Trends in ecology & evolution, 27, 480–488.

Giraldo, N., Salazar, C., Jiggins, C.D., Bermingham, E. & Linares, M. (2008) Two sisters in the same dress: *Heliconius* cryptic species. BMC Evolutionary Biology, 8, 324.

Gómez, A. & Lunt, D.H. (2007) Refugia within refugia: patterns of phylogeographic concordance in the Iberian Peninsula. Phylogeography of southern European refugia (ed. by Weiss, S. & Ferand, N.), pp. 155–188. Springer, Netherlands.

Gustafsson, A.L.S., Skrede, I., Rowe, H.C., Gussarova, G., Borgen, L., Rieseberg, L.H., Brochmann, C. & Parisod, C. (2014) Genetics of cryptic speciation within an arctic mustard, *Draba nivalis*. PloS one, 9(4), e93834.

Harris, H. & Hopkinson, D.A. (1976) Handbook of enzyme electrophoresis in human genetics. North-Holland.

Hebert, P.D.N., Cywinska, A., Ball, S.L. & DeWaard, J.R. (2003) Biological identifications through DNA barcodes. Proceedings of the Royal Society of London, (B) (Biological Sciences), 270, 313–321.

Hewitt, G.M. (1996) Some genetic consequences of ice ages, and their role in divergence and speciation. — Biological Journal of the Linnean Society, 58, 247–276.

Hewitt, G.M. (2011) Mediterranean peninsulas: the evolution of hotspots. Biodiversity hotspots (ed. by Zachos, F. E., & Habel, J. C.), pp. 123–147. Springer, Berlin, Heidelberg.

Hewitt, G.M. (1999) Postglacial re-colonization of European biota. Biological Journal of the Linnean Society, 68, 87–112.

Hinojosa, J.C., Monasterio, Y., Escobés, R., Dincă, V. & Vila, R. (2018) *Erebia epiphron* and *Erebia orientalis*: sibling butterfly species with contrasting histories. Biological Journal of the Linnean Society, 126, 338–348.

Igniatgev, N.N. & Zolotuhin, V.V. (2005) A review of syntomids (Lepidoptera: Syntomidae) of Russia and adjacent territories. Part 1. Genus *Syntomis* Ochsenheimer 1808. Eversmannia, 3/4: 28–54.

Jiggins, C.D. (2017) *The ecology & evolution of* Heliconius *butterflies*. Oxford, University Press.

Knowlton, N. (1993) Sibling species in the sea. Annual review of ecology and systematics, 24, 189–216.

Larkin, M.A., Blackshields, G., Brown, N.P., Chenna, R., McGettigan, P.A., McWilliam, H., Valentin, F., Wallace, I.M., Wilm, A., Lopez, R., Thompson, J.D., Gibson, T.J. & Higgins, D.G. (2007) Clustal W and Clustal X version 2.0. Bioinformatics, 23, 2947–2948.

Lawrence, J.P., Rojas, B., Fouquet, A., Mappes, J., Blanchette, A., Saporito, R.A., Bosque, R.J., Courtois, E.A. & Noonan, B.P. (2019) Weak warning signals can persist in the absence of gene flow. Proceedings of the National Academy of Sciences, 116,19037–19045.

Lees, D.C. & Zilli, A. (2019) Moths: Their biology, diversity and evolution. Natural History Museum, London.

Leimar, O., Tullberg, B.S. & Mallet, J. (2012) Mimicry, saltational evolution, and the crossing of fitness valleys. The Adaptive Landscape in Evolutionary Biology (ed. by Svensson, E., & Calsbeek, R.), pp. 259–270. Oxford University Press, Oxford.

Librado, P., & Rozas, J. (2009) DnaSP v5: a software for comprehensive analysis of DNA polymorphism data. Bioinformatics, 25, 1451–1452.

Malatesta, A. (1985). Geologia e paleobiologia dell’era glaciale. NIS, Roma, 282 pp.

Mayr, E., (1963) Animal species and evolution. Belknap Press of Harvard University Press, Cambridge Mass.

Nei, M. (1978) Estimation of average heterozygosity and genetic distance from a small number of individuals. Genetics, 89, 583–590.

Nguyen, L.T., Schmidt, H.A., von Haeseler, A. & Minh, B.Q. (2014) IQ-TREE: a fast and effective stochastic algorithm for estimating maximum-likelihood phylogenies. Molecular biology and evolution, 32, 268–274.

Obraztsov, N.S. (1966) Die Palaearktischen *Amata* – Arten (Lepidoptera, Ctenuchidae). Veröffentlichungen der Zoologischen Staatssammlung München, 10, 1–383.

Papadopoulou, A., Anastasiou, I. & Vogler, A.P. (2010) Revisiting the insect mitochondrial molecular clock: the mid-Aegean trench calibration. Molecular Biology and Evolution, 27, 1659–1672.

Pardo-Diaz, C., Salazar, C., Baxter, S.W., Merot, C., Figueiredo-Ready, W., Joron, M., McMillan, W.O. & Jiggins, C.D. (2012) Adaptive introgression across species boundaries in *Heliconius* butterflies. PLoS Genetics, 8, e1002752.

Peakall, R.O.D. & Smouse, P.E. (2006) GENALEX 6: genetic analysis in Excel. Population genetic software for teaching and research. Molecular ecology notes, 6, 288–295.

Pérez-Ponce de León, G. & Poulin, R. (2016) Taxonomic distribution of cryptic diversity among metazoans: not so homogeneous after all. Biology letters, 12, 20160371.

Pfennig, D.W. (2012) Mimicry: ecology, evolution, and development. Current Zoology, 58, 603–606.

Pfennig, D.W. & Mullen, S.P. (2010) Mimics without models: Causes and consequences of allopatry in Batesian mimicry. Proceedings of the Royal Society of London, (B), 277, 2577–2585.

Pfenninger, M. & Schwenk, K. (2007) Cryptic animal species are homogeneously distributed among taxa and biogeographical regions. BMC evolutionary biology, 7, 121.

Przybyłowicz, Ł., Lees, D.C., Zenker, M.M., & Wahlberg, N. (2019) Molecular systematics of the arctiine tribe Syntomini (Lepidoptera, Erebidae). Systematic Entomology, 44, 624–637.

Pons, J., Barraclough, T.G., Gomez-Zurita, J., Cardoso, A., Duran, D.P., Hazell, S., Kamoun, S., Sumlin, W.D. & Vogler, A.P. (2006) Sequence-based species delimitation for the DNA taxonomy of undescribed insects. Systematic Biology, 55, 595–609.

Puillandre, N., Lambert, A., Brouillet, S. & Achaz, G. (2012a) ABGD, Automatic Barcode Gap Discovery for primary species delimitation. Molecular ecology, 21, 1864–1877.

Puillandre, N., Modica, M.V., Zhang, Y., Sirovich, L., Boisselier, M.C., Cruaud, C., Holford, M. & Samadi, S. (2012b) Large-scale species delimitation method for hyperdiverse groups. Molecular ecology, 21, 2671–2691.

Racheli, T. & Zilli, A. (1985) Modelli di distribuzione dei Lepidotteri nell’Italia meridionale. Biogeographia, 11, 165–194.

Ragusa, E. (1904) Note Lepidotterologiche. Il Naturalista Siciliano, 17, 18–20.

Reidenbach, K.R., Neafsey, D.E., Costantini, C., Sagnon, N.F., Simard, F., Ragland, G.J., Egan, S.P., Feder, J.L., Muskavitch, M.A.T. & Besansky, N.J. (2012) Patterns of genomic differentiation between ecologically differentiated M and S forms of *Anopheles gambiae* in West and Central Africa. Genome biology and evolution, 4, 1202–1212.

Robuchon, M., Faith, D.P., Julliard, R., Leroy, B., Pellens, R., Robert, A., Thevenin, C., Veron, S. & Pavoine, S. (2019) Species splitting increases estimates of evolutionary history at risk. Biological Conservation, 235, 27–35.

Roca, A.L., Georgiadis, N., Pecon-Slattery, J. & O’Brien, S.J. (2001) Genetic evidence for two species of elephant in Africa. Science, 293, 1473–1477.

Rögl, F. (1999) Mediterranean and Paratethys. Facts and hypotheses of an Oligocene to Miocene paleogeography (short overview). Geologica carpathica, 50, 339–349.

Ruxton, G.D., Sherratt, T.N. & Speed, M.P. (2004) Avoiding attack: the evolutionary ecology of crypsis, warning signals and mimicry. Oxford University Press, Oxford.

Sbordoni, V., Bullini, L., Scarpelli, G., Forestiero, S. & Rampini, M. (1979) Mimicry in the burnet moth *Zygaena ephialthes*: population studies and evidence of a Batesian-Müllerian situation. Ecological Entomology, 4, 83–93.

Sbordoni, V., Bullini, L., Bianco, P., Cianchi, R., De Matthaeis, E. & Forestiero, S. (1982) Evolutionary studies on ctenuchid moths of the genus *Amata*: 2. Temporal isolation and natural hybridization in sympatric populations of *Amata phegea* and *A. ragazzii*. Journal of the Lepidopterists’ Society, 36, 185–191.

Scalercio, S., Cini, A., Menchetti, M., Vodă, R., Bonelli, S., Bordoni, A., Casacci, L., Dinča, V., Balletto, E., Vila, R. & Dapporto, L. (2019) How long are 3 kilometers for a butterfly? Ecological constraints and functional traits explain high genetic differentiation between Sicily and the Italian Peninsula. XXI European Congress of Lepidopterology, University of Molise, Campobasso (Italy), Book of abstracts, 76–77.

Schenk, J.J. (2016) Consequences of secondary calibrations on divergence time estimates. PLoS One, 11(1), e0148228.

Schmitt, T. (2007) Molecular biogeography of Europe: Pleistocene cycles and postglacial trends. Frontiers in zoology, 4, 11.

Schneider, D., Legal, L., Dier,l W. & Wink, M. (1999) Androconial hairbrushes of the *Syntomis (Amata) phegea* (L.) group (Lepidoptera, Ctenuchinae): a synapomorphic character supported by sequence data of the mitochondrial 16S rRNA gene. Zeitschrift für Naturforschung, C, 54, 1119–1139.

Schwarzfeld, M.D. & Sperling, F.A. (2015) Comparison of five methods for delimitating species in *Ophion* Fabricius, a diverse genus of parasitoid wasps (Hymenoptera, Ichneumonidae). Molecular phylogenetics and evolution, 93, 234–248.

Selander, R.K. (1971) Biochemical polymorphism and systematics in the genus *Perumyscus*. 1. Variation in the old field mouse (*Peromyscus polionatus*). University of Texas Publications, 7103, 49–90.

Simon, C., Frati, F., Beckenbach, A., Crespi, B., Liu, H. & Flook, P. (1994) Evolution, weighting, and phylogenetic utility of mitochondrial gene sequences and a compilation of conserved polymerase chain reaction primers. Annals of the entomological Society of America, 87, 651–701.

Stauder, H. (1913) *Syntomis phegea* L. aus dem ӧsterreichischen Litorale und Mitteldalmatien. Zeitschrift für wissenschaftliche Insektenbiologie, 9, 236–239.

Stauder, H. (1928-1929). Genus *Syntomis* O. im zirkum-adriatisch-tyrrhenisch-ligurischen Gebiete. Lepidopterologische Rundschau, 2, (15) 149-154, (16/17) 160-171, (18) 173-176, (19) 187-190, (20) 200-201, (21) 207-210, (22) 215-218, (23) 227-230, (24) 239-242. Entomologische Anzeiger, 9, 10–12.

Struck, T.H., Feder, J.L., Bendiksby, M., Birkeland, S., Cerca, J., Gusarov, V.I., Kistenich, S., Larsson K-H., Liow, L.H., Nowak, M.D., Stedje, B., Bachmann, L. & Dimitrov, D. (2018) Finding evolutionary processes hidden in cryptic species. Trends in Ecology & Evolution, 33, 153–163.

Suc, J.P. & Popescu, S.M. (2005) Pollen records and climatic cycles in the North Mediterranean region since 2.7 Ma. Geological Society, London, Special Publications, 247, 147–158.

Swofford, D.L. & Selander, R.B. (1999) BIOSYS-2. A Computer Program for the Analysis of Allelic Variation in Population Genetics and Biochemical Systematics (Release 2.0). University of Illinois, Urbana-Champaign.

Taberlet, P., Fumagalli, L., Wust-Saucy, A. & Cosson, J. (1998) Comparative phylogeography and postglacial colonization routes in Europe. Molecular Ecology, 8, 1923–1934.

Tamura, K., Stecher, G., Peterson, D., Filipski, A. & Kumar, S.. (2013). MEGA6: molecular evolutionary genetics analysis version 6.0. Molecular biology and evolution, 30, 2725–2729.

Turati, E. (1917) Revisione delle *Syntomis* Paleartiche a doppio cingolo giallo, e saggio di una classificazione delle varie forme e specie. Atti della Società italiana di Scienze naturali, 56, 179–232, pls 2–8.

Verity, R. (1914) Contributo allo studio della variazione nei Lepidotteri tratto principalmente da materiale di Toscana, delle Marche e di Calabria. Bollettino della Società Entomologica Italiana, 45, 203–238, 1 pl.

Wiens, J.J. (2007) Species delimitation: new approaches for discovering diversity. Systematic Biology, 56, 875–878.

Wiens, J.J. & Servedio, M.R. (2000) Species delimitation in systematics: inferring diagnostic differences between species. Proceedings of the Royal Society of London,(B), 267, 631–636.

Zhang, J., Kapli, P. & Stamatakis, A. (2013) A general species delimitation method with applications to phylogenetic placements. Bioinformatics, 29, 2869–2876.

Zilli, A. (1996) Colour polymorphism of *Callimorpha dominula* (Linnaeus, 1758) in Italy, and the problem of polytopic subspecies (Lepidoptera, Arctiidae, Callimorphinae). Mitteilungen der Münchner Entomologischen Gesellschaft, 86, 79–98.

